# Uncover spatially informed shared variations for single-cell spatial transcriptomics with STew

**DOI:** 10.1101/2023.10.10.561789

**Authors:** Nanxi Guo, Juan Vargas, Douglas Fritz, Revanth Krishna, Fan Zhang

## Abstract

**Motivation:** The recent spatial transcriptomics (ST) technologies have enabled characterization of gene expression patterns and spatial information, advancing our understanding of cell lineages within diseased tissues. Several analytical approaches have been proposed for ST data, but effectively utilizing spatial information to unveil the shared variation with gene expression remains a challenge.

**Results:** We introduce STew, a Spatial Transcriptomic multi-viEW representative learning method, to jointly analyze spatial information and gene expression in a scalable manner, followed by a data-driven statistical framework to measure the goodness of model fit. Through benchmarking using Human DLPFC data with true manual annotations, STew achieved superior performance in both clustering accuracy and continuity of identified spatial domains compared with other methods. STew is also robust to generate consistent results insensitive to model parameters, including sparsity constraints. We next applied STew to various ST data acquired from 10x Visium and Slide-seqV2, encompassing samples from both mouse and human brain, which revealed spatially informed cell type clusters. We further identified a pro-inflammatory fibroblast spatial niche using ST data from psoriatic skins. Hence, STew is a generalized method to identify both spatially informed clusters and disease-relevant niches in complex tissues.

**Availability:** Source code and the R software tool STew are available from github.com/fanzhanglab/STew.

**Supplementary information:** Supplementary data are provided.

## Introduction

Single-cell transcriptomics are revolutionizing our comprehension of transcriptional heterogeneity for complex diseases, such as tumors, autoimmune disorders, neurology diseases. These ailments usually attack human tissues and organs, often stemming from various immune dysfunctions and genetic dysregulation (Shalek and Benson 2017; Stubbington *et al*. 2017; Papalexi and Satija 2018). Recent advancements in single-cell multimodal technologies allow for the simultaneous measurement of multiple data modalities, thereby enhancing our ability to categorize diseased tissues more precisely. For instance, CITE-seq, quantifying both transcriptomics and proteomics, has been used to identify pathogenic cell types in disease contexts, including our recent single-cell multi-modal cross-dataset integration work on inflammatory disorders (Korsunsky *et al*. 2019; Zhang *et al*. 2019, 2022; Kang *et al*. 2021; Reshef *et al*. 2021). However, pinpointing the exact locations where the identified key cell phenotypes drive tissue inflammation remains largely unclear. The recent emergence of spatial transcriptomics (ST) technologies has bridged this gap, facilitating concurrent measurements of gene expression with spatial localization, preserving the spatial information through the analysis of the intact tissue samples (Larsson, Frisén and Lundeberg 2021; Moses and Pachter 2022). This spatial awareness provides a deeper understanding of the intricate relationships between the cells and their neighbors, unlocking new insights into the functions and mechanisms of cellular phenotypes, and thereby catalyzing the progression of innovative therapeutic strategy development.

Multiple cutting-edge ST technologies have been commercialized, varying significantly in aspects such as resolution and throughput (Larsson, Frisén and Lundeberg 2021; Moses and Pachter 2022; Williams *et al*. 2022; Vandereyken *et al*. 2023). For example, the 10x Genomics Visium has the capability to analyze tens of thousands of genes on thousands of locations by spatial barcoding at 55-μm resolution (Ståhl *et al*. 2016). In parallel, Slide-seq (Rodriques *et al*. 2019) employs arrays comprised of 10-μm barcoded beads followed by an improved version Slide-seq V2 (Stickels *et al*. 2020). In situ sequencing (ISS) represents another approach that could measure the entire transcriptome at a single-cell resolution (Ke *et al*. 2013), such as the newly launched Xenium platform(Janesick *et al*. 2022). Moreover, MERFISH, a technology grounded in single-molecule fluorescent in situ hybridization (smFISH), demonstrates the capacity to detect hundreds to tens of thousands of genes at the subcellular level (Chen *et al*. 2015; Moffitt *et al*. 2016).

To overcome current technological constraints, researchers are developing robust and sensitive computational algorithms primarily aimed at addressing the questions of transcript distribution prediction and cell type deconvolution. A recent comprehensive benchmarking study revealed that tools such as Tangram (Biancalani *et al*. 2021), gimVI (Lopez *et al*. 2019), and SpaGE (Abdelaal *et al*. 2020) have proven adept at predicting the spatial distribution of RNA transcripts. Simultaneously, Cell2location (Kleshchevnikov *et al*. 2022) and RCTD (Cable *et al*. 2021) have emerged as relatively potent methods for deconvoluting cell types in spots (Li *et al*. 2022). Looking ahead, it is anticipated that the challenges surrounding cell type deconvolution will diminish in the near future, propelled by swift advancements that are amplifying the resolution of ST technologies.

In this study, we delve into the advantages of leveraging spatial information to unveil the shared variation in gene expression space through the lens of representative learning. While several methods have been developed, it remains a computational challenge given the complexity of the disease tissue and the challenges of inferring biologically meaningful phenotypes. For example, stLearn (Pham *et al*. 2020) introduces a normalization method that incorporates spatial location and tissue morphology to adjust gene expression values. However, stLearn requires the availability of H&E image; in their absence, it resorts to employing single model-based dimensionality reduction methods such as principal component analysis (PCA) and Louvain clustering. MERINGUE constructs a weighted graph that combines spatial and transcriptional similarities (Miller *et al*. 2021). This approach exploits spatial cross-correlation with gene expression patterns, restricting inferred interactions to short-range regions only. Furthermore, SpaceFlow (Ren *et al*. 2022) amalgamates the pseudotime concept with the spatial locations of cells using spatially regularized deep graph networks to elucidate cellular spatiotemporal patterns. To address this critical problem, we present STew, a Spatial Transcriptomic multi-viEW representation learning method grounded in sparse canonical correlation analysis and graph representation to jointly analyze spatial information and gene expression in a scalable fashion. Within STew, we have incorporated a data-driven statistical framework to assess the goodness of fit for biological axes-specific genes. STew outputs distinct spatially informed cell gradients, robust spatially informed clusters, and statistical goodness of model fit to uncover significant genes that reflect subtle spatial changes. To validate its efficacy, we benchmarked STew by applying it to four ST datasets, encompassing a range of healthy and diseased tissues analyzed using distinct technologies.

## Methods

### Dataset

We analyzed four single-cell spatial transcriptomic datasets, which are available from their original publications. For the 10x Visium human dorsolateral prefrontal cortex (DLPFC) data, raw count matrix, histology image, and spatial data are accessible in the “spatialLIBD package” (https://research.libd.org/spatialLIBD/) (Maynard *et al*. 2021; Pardo *et al*. 2022). Manual annotation information for this dataset is also provided. The DLPFC data has been served as a gold standard to test the performance of computational methods for spatial transcriptomics data analyses. We also analyzed the Mouse Brain Anterior dataset which was obtained from https://satijalab.org/seurat/articles/spatial_vignette.html. The Slide-seqV2 mouse hippocampus dataset was accessible from the Broad Institute database “Single Cell Portal” (https://singlecell.broadinstitute.org/single_cell/study/SCP815/highly-sensitive-spatial-transcriptomics-at-near-cellular-resolution-with-slide-seqv2#study-download) (Stickels *et al*. 2020). Further, we focus on analyzing the single-cell spatial transcriptomics data generated from the psoriasis skin, a heterogeneous disease tissue affected by inflammatory disorders.

This dataset is accessible at GEO GSE173706 and GSE225475 (Ma *et al*. 2023). Furthermore, we benchmarked our results with other state-of-the-art methods, including SpaceFlow V1.0.4, stLearn V0.4.12, and Seurat V5 with default parameters.

### Overview of STew

We describe the methodology of STew as follows (**Fig. 1**). The main idea of STew is taking advantage of both spatial information and gene expression data by extracting shared information through multi-view representative learning. STew incorporates the graph-embedded information into the sparse canonical correlation analysis framework by explicitly modeling the sparsities to capture the maximal covariance.

**Fig. 1.**
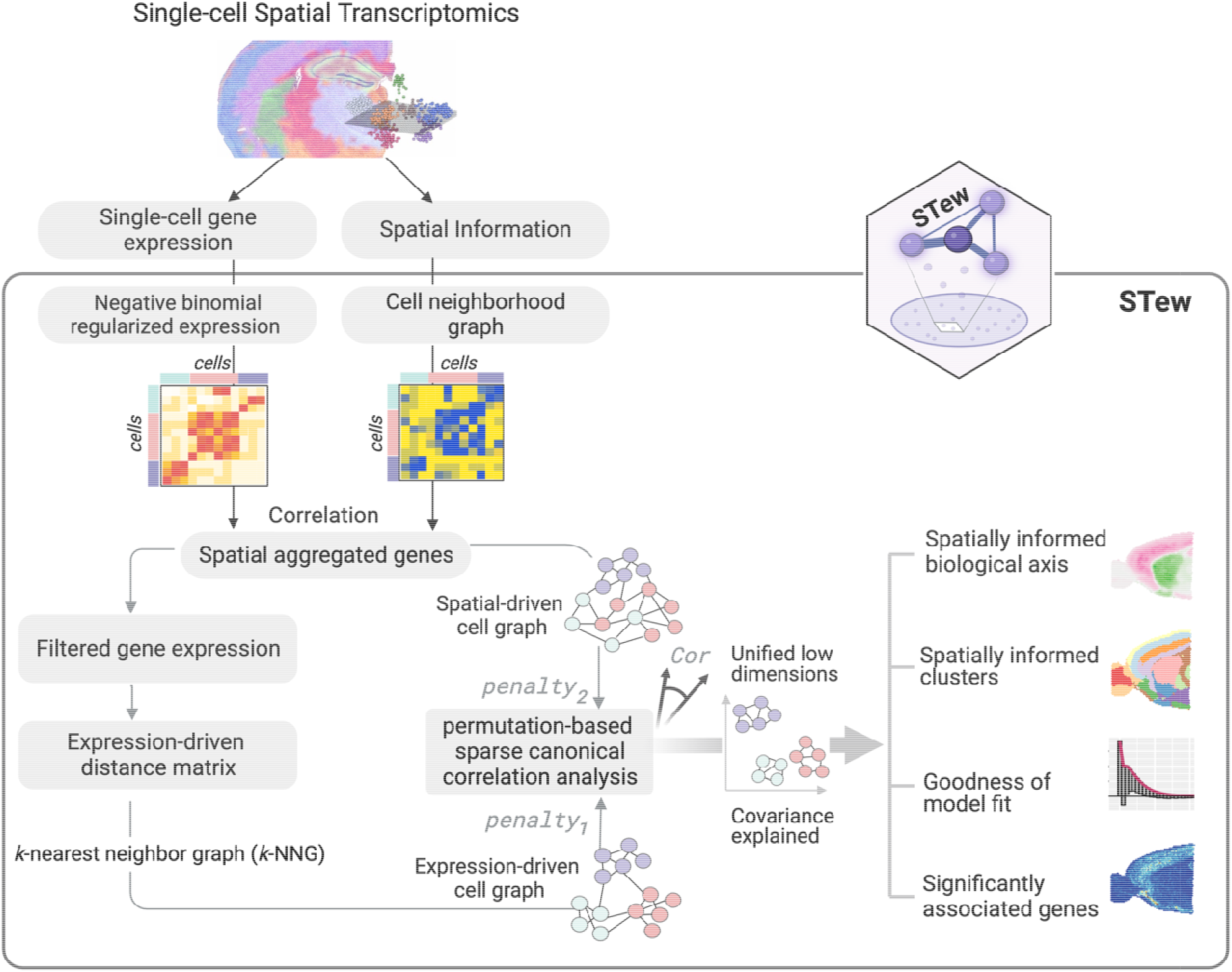
Overview of STew methodology.

We consider a spatial transcriptomic data as (*Y, Z*) ∈ *R*^*n*×*p*^ × *R*^*n*×*2*^ which denotes two views of collected data points on the same set of *n* observations, where *p* represents the number of features. *Y ∈ R*^*n*×*p*^ represents a gene expression matrix with *n* cell spots that measures *p* genes, and *Z ∈ R*^*n*×2^ represents the *n* spots that are distributed on the 2-dim coordinates. For the spatial positional coordinates of the spots, STew characterizes cell-cell adjacency relationships by assigning binarized weight to each pair of cells using Delaunay triangulation, resulting in a binary adjacency matrix *M ∈ R*^*n*×*n*^ as a spatial-driven cell graph, where:

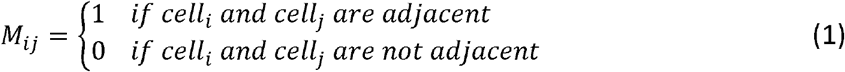

For the gene expression data *Y*, STew utilizes a regularized negative binomial regression model to normalize gene expression data to control for variation that may arise due to differences in sequencing depth or other potential technical inconsistencies. STew defines spatially auto-correlated genes when their expressions vary across different regions of a tissue, often clustering in functionally related groups specific to that tissue region. STew detects such spatially aggregated genes by computing Moran’s I (Miller *et al*. 2021):

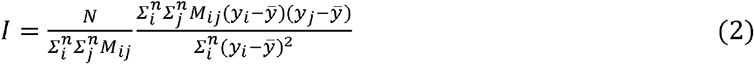

Where *y*_*i*_ denotes a normalized gene expression vector of gene *i* in *Y*. The adjacency matrix describes positive spatial autocorrelations across *n* cells. The Benjamini-Hochberg procedure is used to correct for multiple testing and control for false discovery.

While global measure of spatial association (Moran’s I) provides a single value summarizing the overall spatial pattern of the gene expression, STew next filters out spatial patterns driven by neighborhood of cell spot *i*: small numbers of cells by Local Indicator of Spatial Association (LISA) (Anselin 2010). For each neighborhood of cell spot *i*:

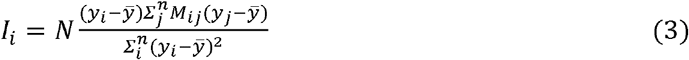

which is related to Moran’s I by:

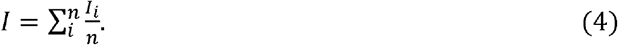

Specifically, STew further determines the spatial pattern scale based on the percentage of cells with a statistically significant LISA value. As a default setting, STew focuses on spatially heterogeneous genes with a p-value below the 0.05 alpha threshold that are driven by at least 5% of the cells. After such steps, we obtain an updated gene expression matrix, denoted as *Y ∈ R*^*n*×q^, where *q* represents the number of spatially aggregated genes.

Next, STew characterizes the cell-cell similarity matrix for *Y*_*nxq*_ by building a weighted *k*-nearest node and its *k*-nearest neighbors based on the Euclidean distance in terms of the gene neighbors’ graph (K-NNG), where each node represents a cell, and an edge is drawn between a node and its *k*-nearest neighbors based on the Euclidean distance in terms of the gene expression. Users may adjust the value of *k* to define the number of neighbors each cell is connected to, 200 by default.

The Euclidean distance is given by:

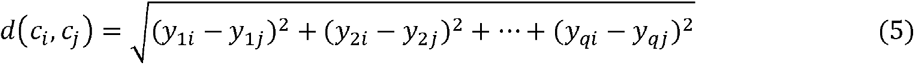

where *c* _*i*_ and *c* _*j*_ denote a pair of cells, and *d(c* _*i*,_ *c* _*j*_ ) is the Euclidean distance between cells in terms of *q* gene expression vectors.

The weighted KNN graph is given by:

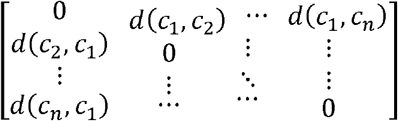

The weighted KNN graph then be converted into a binary adjacency matrix, denoted as *V ∈ R*^*n*×*n*^:

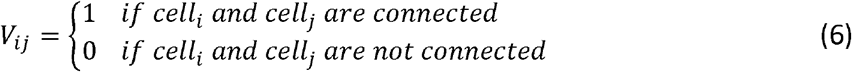

where *j* is one of the *k* nearest neighbors of *i*.

In summary, this first step captures the relationships between cells in the high-dimensional space by converting the original gene expression into gene expression-driven adjacency matrix, denoted as *V ∈ R*^*n*×*n*^, and creating a spatial-driven cell neighborhood graph, denoted as *M ∈ R*^*n*×*n*^. These two adjacency matrices will be the inputs for the following step.

Considering the different levels of sparsity and complexity of the two views, we incorporated penalized canonical coefficients by applying *L*_1_-norm penalties to the coefficients in Equation covariance matrices Σ_11_ and Σ_22_ are usually replaced by identity matrix *I* to obtain decent (7). When dealing with high-dimensional problems, it has been approved that the within-set covariance matrices Σ_11_ and Σ_22_ are usually replaced by identity matrix / to obtain decent results (Witten, Tibshirani and Hastie 2009). Thus, we seek new pairs of sparse canonical coefficients *w*_1_ ∈ *R*^*n*×1^ and *w*_2_ ∈ *R*^*n*×1^ for *M* and *V*, respectively, by maximizing the correlation between (*Mw*_1_, *Vw*_2_) through this optimization problem:

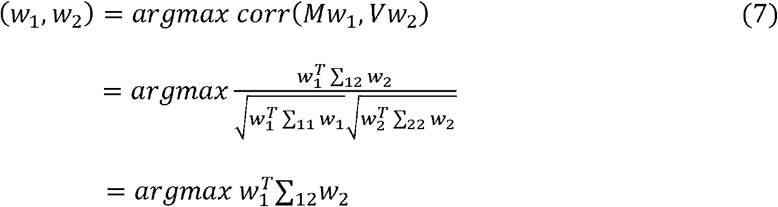

Subject to 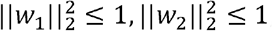, ‖ *w*_1_ ‖ _1_ ≤ *c*_1_, *w*_2_ ‖ _1_ ≤ *c*_2_

Here, 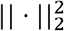 indicates the squared Frobenius norm (the sum of squared elements of the matrix), and 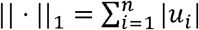 is known as Lasso. In ST data, we usually have a greater number of detected genes than that of spots, which means that these inverses of sample covariance matrices do not exist. Thus, we apply regularizations in the optimization.

In summary, this procedure yields sparse vectors *w*_1_ and *w*_2_ for *c*_1_ and *c*_1_ chosen appropriately. Here, we restrict *c*_1_ and *c*_*2*_ to the range of 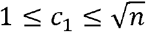 and 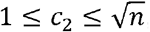, by scaling the *L*_1_ regularization term with respect to *M* and *V*, resulting in penalty 0 < λ_1_ ≤ 1, and 0 < λ_1_ ≤ 2. Larger *L*_1_ bound corresponds to less penalization.

The values of *c*_1_ and *c*_2_ are determined through a parallelized permutation testing approach. For each candidate penalty parameter within the range of (0, 1), we randomly permute the rows of matrices *M* and *V k* times to generate 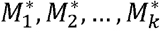 and 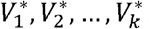. Then, we run sparse CCA on each permuted dataset 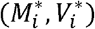 to obtain factors (*w*_1*i*_, *w*_2*i*_). We then compute the correlation *C*_*i*_ *= corr (M*_*i*_*w*_1*i*_, *V*_i_ *w*_2*i*_) and *C* = *corr* (*Mw*_*1*_, *Vw*_2_), where (*M, V*) is from the original data and (*w*_1_,*w*_2_) is the obtained factors from the original data. Lastly, we use Fisher’s transformation to convert the correlation to a random variable that is approximately normally distributed. Let *Fisher*(*c*) denotes the Fisher transformation of *c*, and then compute a z-statistic for *Fisher*(*C*) using 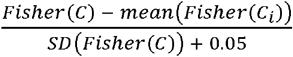. Here, 0.05 is added to the denominator to avoid division by zero issues. The magnitude of the z-statistic serves as an indicator of the significance of the corresponding tuning parameter value for matrices *M* and *V*. A larger absolute value of the z-statistic suggests a superior fit of the sparse CCA model to the data, given that specific combination of tuning parameters. This implies a reliable and meaningful association between the matrices under consideration.

As outputs, we generate low-dimensional embeddings, termed as sparse canonical vectors (sCVs), that reflect the shared covariance. Furthermore, we detect cell type clusters within a tissue via smart local moving (SLM) algorithm by constructing a shared neighborhood graph (SNN) based on sCVs. Other downstream analyses such as data visualization can be performed on these derived sCVs.

### Statistical count modeling and goodness of model fit evaluation

To determine the transcripts that are significantly associated with the sparse canonical vectors or identified clusters, we should select the right statistical model for testing given the variety of means and standard deviation of genes of interest. Thus, we incorporate statistical modeling comparison into STew by evaluating various count models to deal with the overdispersion attribute, including zero-inflated Poisson, negative binomial, zero-inflated negative binomial, Poisson Generalized Linear Model (GLM), and hurdle, applying on raw counts instead of transformed values. We assess model performance using Akaike Information Criterion (AIC) and Rootograms (Kleiber and Zeileis 2016) to determine the best-fitting model considering the issues of over-estimate or under-estimate. In addition, we evaluate the mean and variation captured under the gene expression space for each tested count model. Our framework also accounts for covariates in the Likelihood Ratio Test (LRT) by comparing the full model with the null model with a significance level of 0.05. We then generate coefficient and p-value for each tested gene, and report statistically significant genes as outputs.

### Performance Evaluation

To evaluate the performance of the clustering results, we measured the similarities of identified clusters for STew and other benchmarking methods with the true manual annotations using ARI (adjusted rand index):

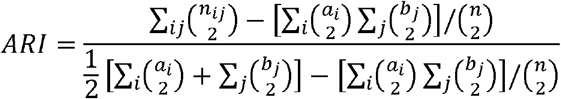

Where *a*_*i*_ = Σ_*j*_ *n*_*ij*_, *b*_*j*_ = Σ_*i*_ n_*ij*_, and here n_*ij*_ represents the number of agreements between true manual annotations and predicted cluster labels.

Further, we quantified the continuity and smoothness of the identified spatial domains by calculating a percentage of abnormal spots (PAS) score. As the PAS score is calculated based on the proportion of spots with a cluster label that is different from at least six of its neighboring ten spots (Shang and Zhou 2022), lower PAS score indicates better spatial continuity and smoothness of in the identified cluster.

## Results

### STew robustly recapitulates true manual annotations for human brain ST data

We applied STew on human dorsolateral prefrontal cortex (DLPFC) data, which measured 33,538 genes on 3,639 cell spots, generated from 10x Visium (Maynard *et al*. 2021). We first identified spatially informed sparse canonical vectors (sCVs) and then performed clustering on them. In the STew framework, we automatically identified the optimal penalty parameters through cross-validation, measured the correlations between different sCVs, and identified the correlations that were reflected by each of the sCVs (**Fig. 2A-D**). Further, we estimated the optimal penalty parameters for sparse canonical correlation by comparing the clustering results with the true manual annotations using ARI (**Fig. 2A**), which suggests that the accuracies of clustering results are relatively robust but vary slightly according to the penalties (**Supplementary Figure 1**). The variations among top 20 sCVs are orthogonal to each other (**Fig. 2B**), and each of the low-dimensional embeddings captures different shared variation between the gene expression and spatial regions (**Fig. 2C**). Consistent with these, we visualized each sCV at the spatial image, and different embeddings reflect distinct spatial region relevant niches (**Fig. 2D**).

**Fig. 2.**
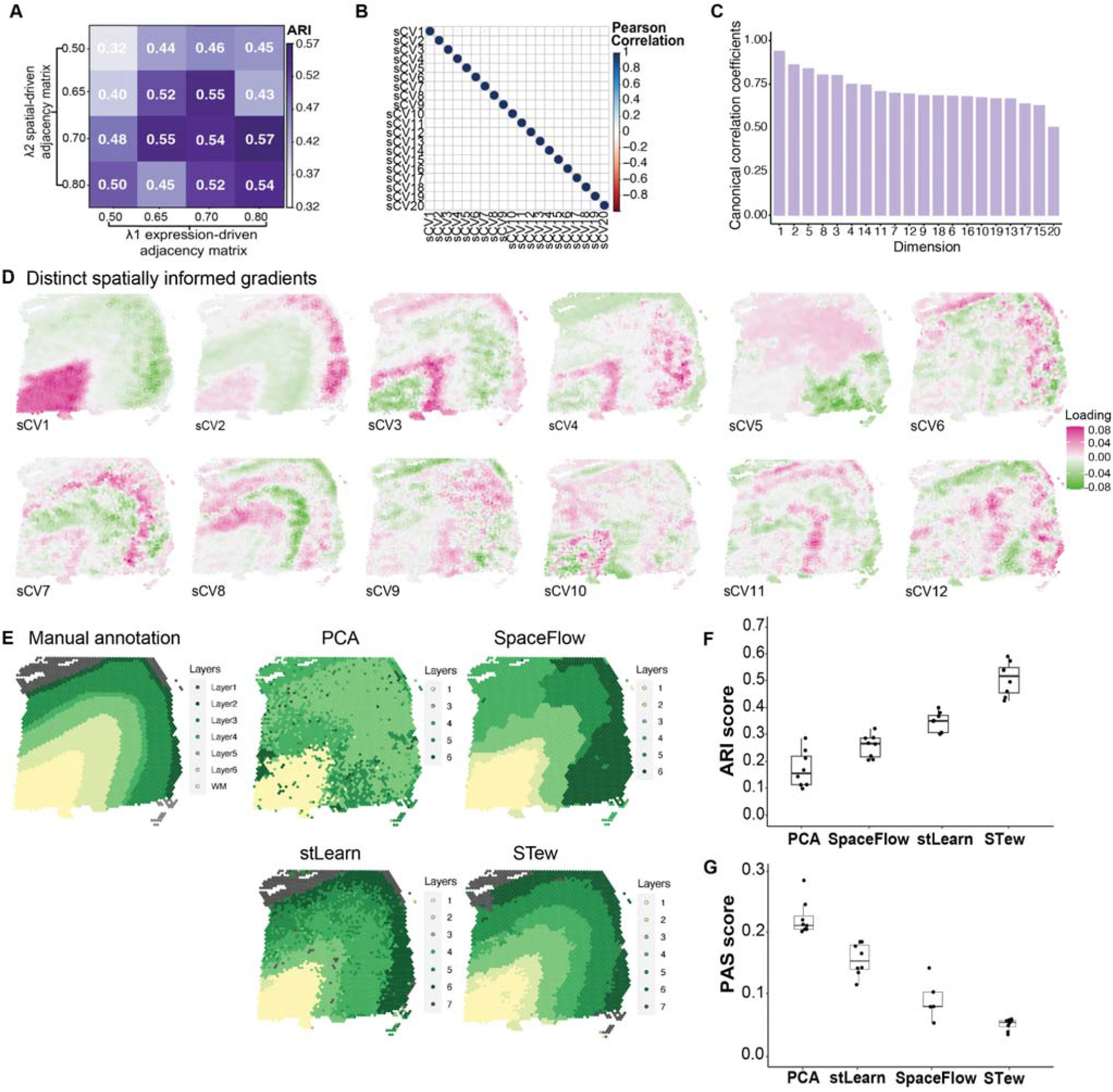
Benchmarking STew on 10x Visum dorsolateral prefrontal cortex data from DLPFC. **A**, ARI (Adjusted Rand Index) scores to evaluate clustering accuracy by comparing with manual annotations. We tested different sparsity constraints in STew regarding their performance. **B**, Pearson correlation coefficients of top 20 sCVs. **C**, Barplot of canonical correlation coefficients of top 20 sCVs. **D**, Distinct spatially informed gradients. **E**, Manual annotations of DLPFC in Slide 151673 and cell type clusters identified by PCA, SpaceFlow, stLearn, and STew using default parameters. **F**, Clustering accuracy in recapitulating the true manual labels measured by ARI, with different numbers of low dimensions in each of the algorithms, in particular top 6, 8, 10, 12, 14, 16, 18, and 20 components. In the boxplot, the center line, box limits and whiskers denote the median, upper, and lower quartiles, and 1.5 interquartile range, respectively. **G**, Percentage of abnormal spots (PAS), the proportion of spots with a cluster label that is different from at least six of its neighboring ten spots.

We further benchmarked the performance of STew with other state-of-the-art methods. By comparing STew-derived clusters with the true manual annotations using ARI (adjusted rand index), we found that STew (median ARI=0.52) outperformed PCA (median ARI=0.16), SpaceFlow (Ren *et al*. 2022) (median ARI=0.27), and stLearn (median ARI=0.35) regarding clustering accuracy (**Fig. 2E, F**). In addition, we measured the continuity and smoothness of the identified spatial domains by percentage of abnormal spots (PAS) score, which reviews that STew (median PAS=0.06) is smoother and more continuous than PCA (median PAS=0.21), stLearn (median PAS=0.16), and SpaceFlow (median PAS=0.08) (**Fig. 2G**). Lower PAS score here indicates a spot homogeneity within the identified clusters. After testing the effects of using different numbers of low-dimensional embeddings as inputs, we note that all the involved methods achieved consistent performances reflected by the ARI (**Fig. 2F**); STew achieves slightly better consistency than other methods regarding the smoothness of the identified spatial domains (**Fig. 2G**).

### STew is scalable to automatically determine the best model of fit for significant gene identification on mouse brain datasets

To evaluate the scalability of STew, we first applied STew on a mouse brain spatial transcriptomic dataset generated from Slide-seqV2, almost single-cell resolution. For this mouse hippocampus data, it measured on 41,786 locations generated from Slide-seqV2 (Stickels *et al*. 2020). We clustered on the spatially informed low dimensions, top 20 sCVs, to identify different layers of mouse brain clusters (**Fig. 3A, Supplementary Figure 2**). The biologically meaningful markers clearly reflect the regional dissociation of the tissue and are consistent with the known anatomic structures (**Fig. 3B**) (Stickels *et al*. 2020). Next, we benchmarked five different statistical models with the potential to model overdispersed count data, aiming to pinpoint the right model for identifying significant genes associated with the discerned spatial informed axes or clusters.

**Fig. 3.**
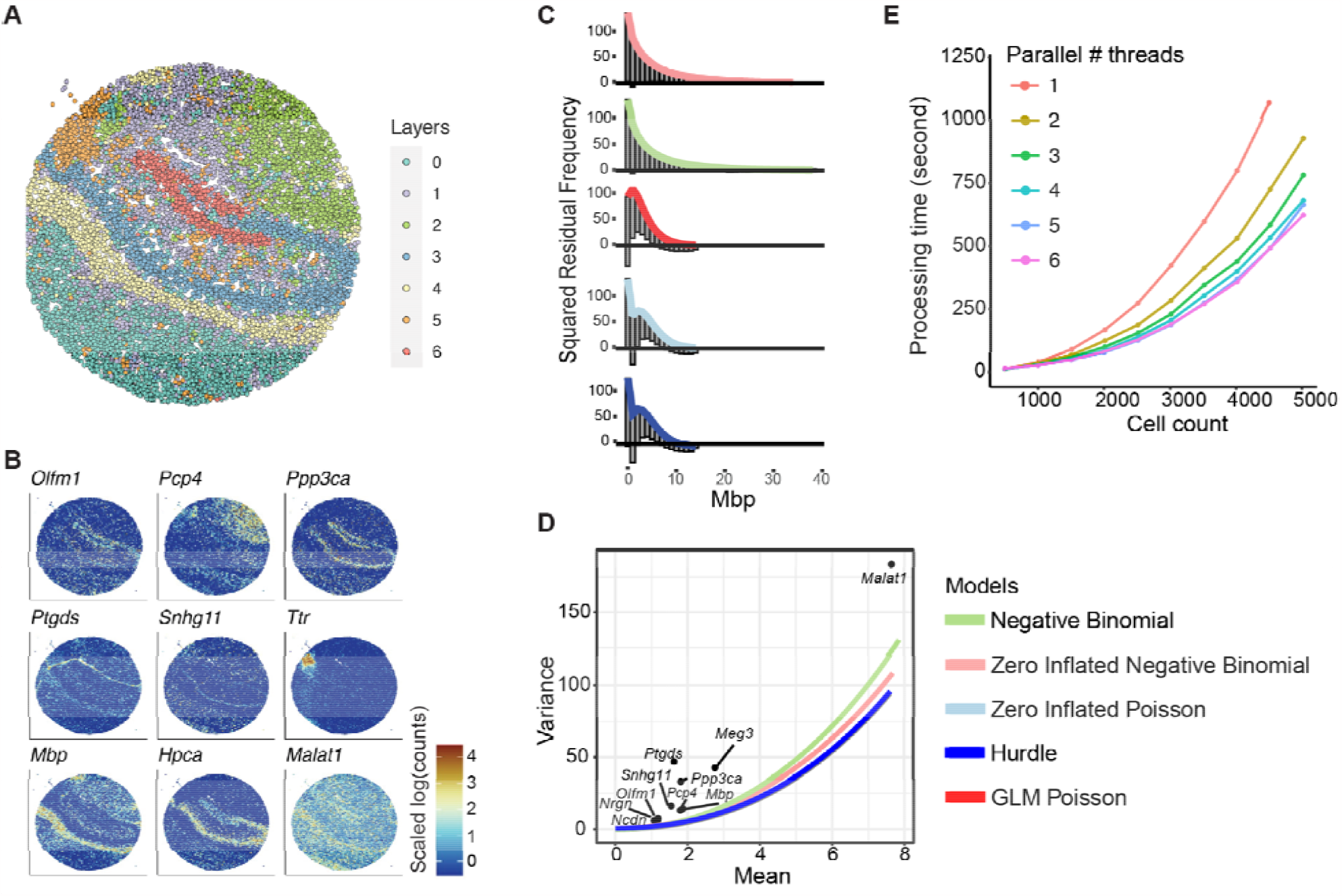
Applying STew on mouse hippocampus data from Slide-seqV2. **A**, Identified cell type clusters of mouse hippocampus region on data from Slide-seqV2. **B**, Gene expression distributions in the spatial domains that reflected spatial region segregation. **C**, Rootogram that measures squared residual frequency for each tested model. X-axis is the counts for gene *Mbp*. **D**, Benchmarking five statistical models that capture mean and variance for each gene. Colors indicate different distributions. Note that the Zero inflated Poisson and GLM Poisson are overlapped with hurdle distribution. **E**, Scalability of parallel computing of STew regarding cell count and # threads on 15 genes.

Interestingly, both negative binomial and zero-inflated negative binomial models exhibited superior performance compared to the other models, as indicated by Rootogram analysis to assess the goodness of model fit at the count level (**Fig. 3C**). Particularly, the Rootogram quantified squared residual frequency for gene *Mbp* (myelin basic proteins) counts, a major constituent of the myelin sheath of oligodendrocytes and Schwann cells in the nervous system.

We observed that negative binomial and zero-inflated negative binomial models fitted the expression distribution very well, but GLM Poisson, zero-inflated Poisson, and Hurdle underestimated the zero counts and over-estimated the low counts (**Fig. 3C, Supplementary Fig. 3**). Further, we benchmarked these statistical count models by evaluating the mean and variance captured for selected genes across a range of counts (**Fig. 3D**). Ultimately, we have incorporated this rigorous statistical model benchmarking within the STew framework, empowering users to automatically select the most appropriate model for analyzing spatial transcriptomic features, which exhibits distributions usually depending on sequencing depth and technological resolution.

Another significant contribution of STew over traditional canonical correlation analysis is the incorporation of parallel computing for the integration step, catering specifically to the analysis of large ST datasets. Given that the number of captured cells may vary depending on the specific tissue context and the technology utilized, this implementation is critical. By operating on different parallel threads, we are able to evaluate the scalability of STew on mouse hippocampus data by the random downsampling of selected numbers of cell spots (**Fig. 3E**). These analyses affirm that STew is scalable to large ST datasets, thereby facilitating the identification of spatially informed niches in complex tissue structures.

Next, we applied STew on a mouse brain anterior dataset acquired by 10x Visium which contains 31,053 genes and 2,696 locations, revealing spatially informed clusters (**Fig. 4A**) and distinct biologically meaningful gradients that capture different layers in the mouse brain (**Fig. 4B, Supplementary Figure 4**). Taking sCV1 as an example, we presented the statistics generated from the zero-inflated negative binomial modeling. The genes with positive estimates, which are markers for the cells with positive loadings in sCV1, like gene *Cabp1* (**Fig. 4C, D**).

**Fig. 4.**
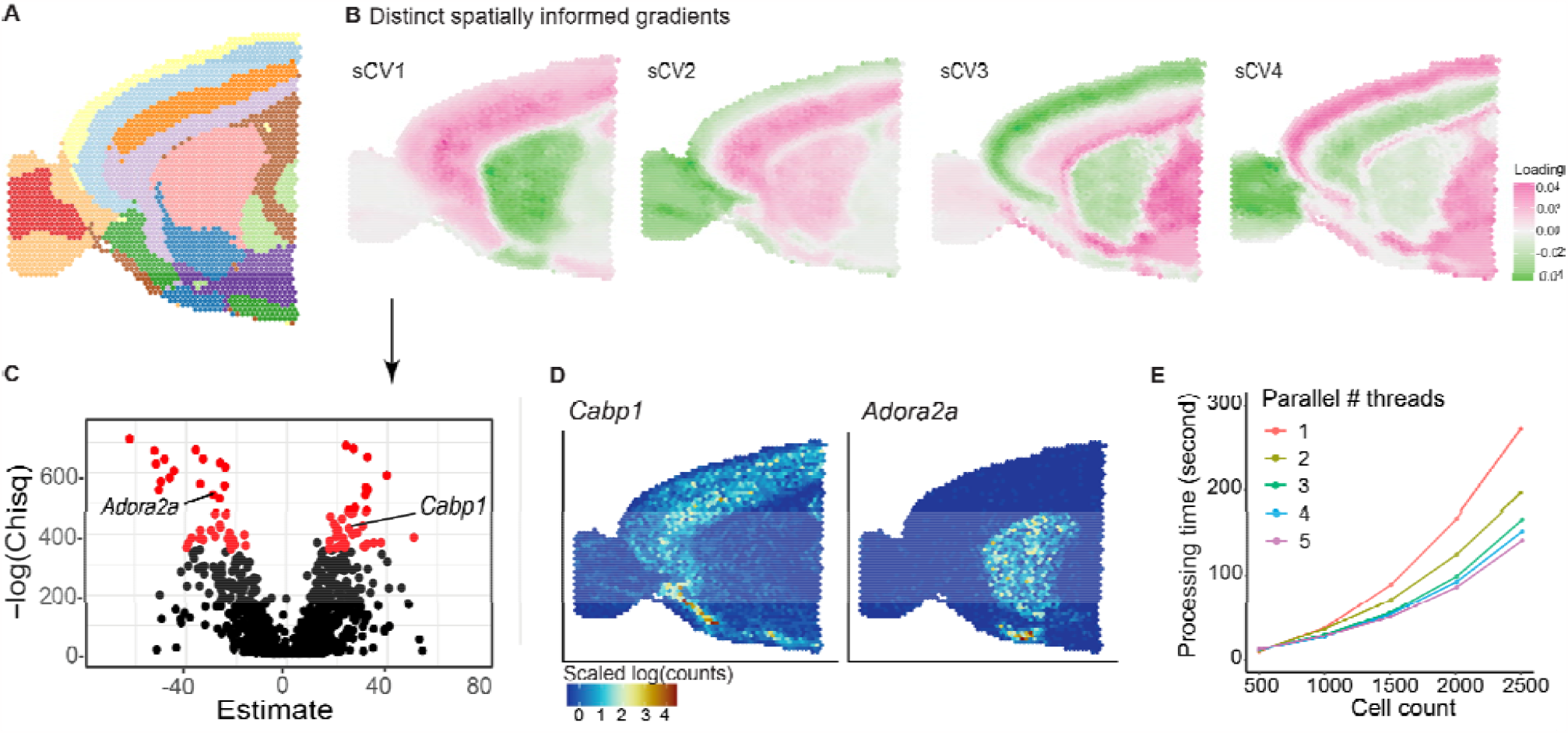
Applying STew on mouse brain data from 10x Visum. **A**, Cell type clusters of 10x Visium Mouse Brain Anterior 1 using STew. **B**, Top spatially informed gradients reflect the spatial domain variation. **C**, Statistics of sCV1 associated genes based on Zero-inflated negative binomial modeling. Red color indicates statistically significant genes. **D**, Gene expression pattern on the image for the selected genes that are significantly associated with sCV1. **E**, Scalability of parallel computing of STew regarding cell count change and number of threads on 600 highly variable genes.

Similarly, genes with negative estimates indicate markers for cells with negative loadings in sCV1. For example, gene *Adora2a* (Adenosine A2a Receptor) (**Fig. 4C, D**), an adenosine receptor group of G-protein-coupled receptors controlling synaptic plasticity, plays a critical role in modulating anxiety and sleep (Hohoff *et al*. 2020). We further evaluated the scalability of STew on the top 600 highly variable genes, which confirmed that STew is able to generate spatially aware results in 5 minutes using a single computing thread and around 3 minutes using two parallel threads (**Fig. 4E**).

### STew characterizes the spatial organization of heterogeneous cell types in psoriatic skin

Inflammatory diseases affect different human tissues which are usually more heterogeneous and complex than the well-structured human brain. To evaluate the sensitivity of STew, we applied it to analyze a recent spatial transcriptomic dataset generated from psoriatic skin using 10x Genomics (Ma *et al*. 2023). We identified five major cell types, including keratinocyte, smooth muscle, fibroblast, eccrine gland, and myeloid and T cells (**Fig. 5A**). Our annotations of these cell types are based on the canonical marker genes reported from the original study (Ma *et al*. 2023). For instance, we are able to allocate smooth muscle cells, which aligned with corresponding marker gene expressions of *DES* (Pearson r = 0.18, *p* = 3.4e-11), *ACTG2* (Pearson r = 0.19, *p* = 1.2e-12), *TAGLN* (Pearson r = 0.33, *p* = 2.2e-16), and *MYH11* (Pearson r = 0.14, *p* = 9.5e-8) (**Fig. 5B**). In addition, we detected eccrine gland cells which were formed by two distant regions, but they consistently expressed the same marker genes, including *PIP, MUCL1, SCGB1B2P*, and *SCGB1D2* (**Fig. 5C**). We also identified fibroblasts that highly expressed marker genes such as *COL1A1, DCN, CFD*, and *COL3A1* (**Fig. 5D**). Note that, fibroblasts, smooth muscle, and eccrine gland cells are very heterogeneous and mixed in the diseased skins, which are typically situated deeper within the dermis. This complexity renders them difficult to distinguish using solely gene expression patterns, since the spatial location of these tissue-specific cell types can create unique microenvironments or niches. Our algorithm is able to delineate these cells into continuous and smooth spatial domains measured by a PAS value of 0.084, and further maintain the concordance with cell type lineage marker expressions (**Fig. 5 A-D**). This analysis demonstrates the capability of STew in identifying the shared variations from both gene expression and anatomical spatial domains within the context of disease.

**Fig. 5.**
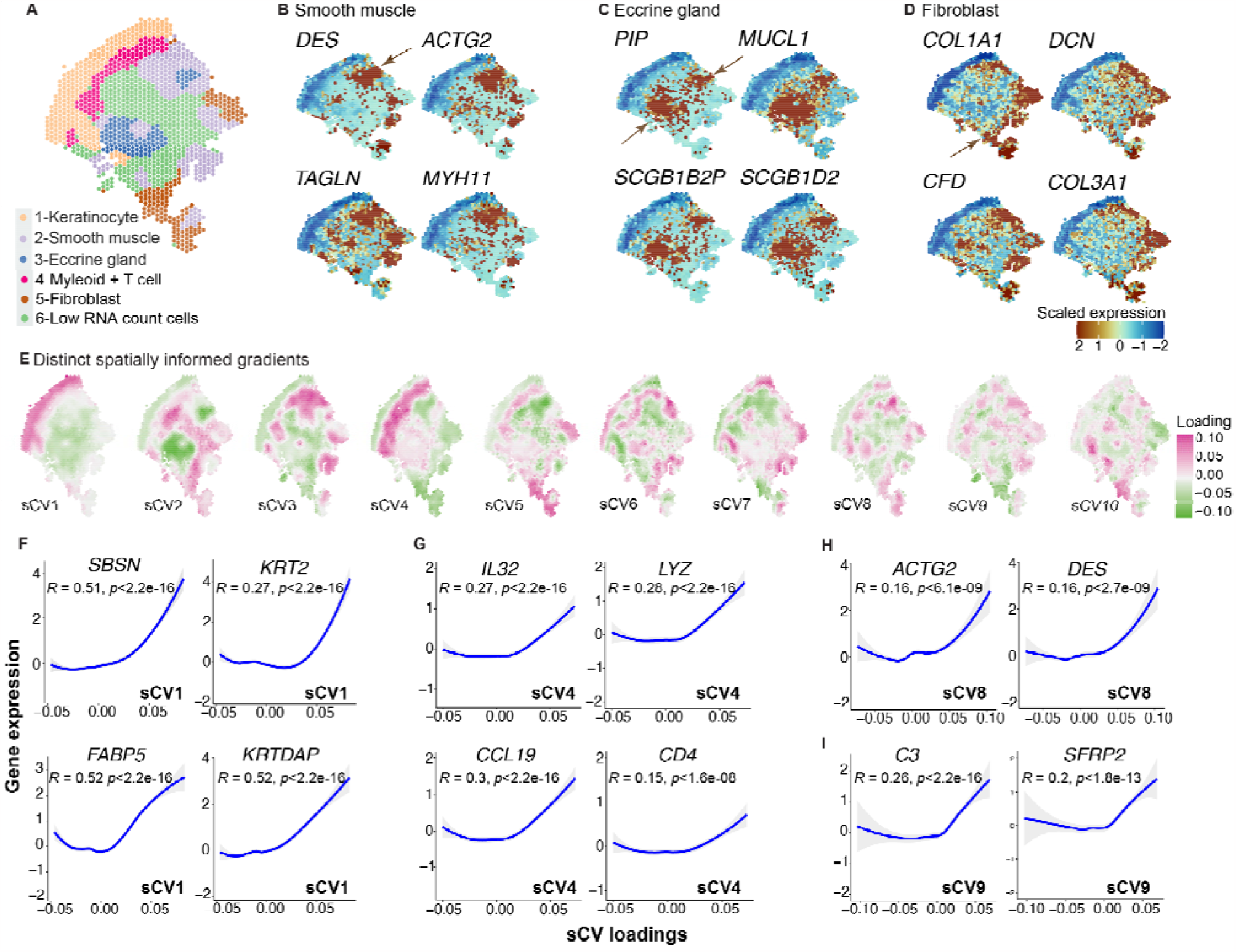
Analysis of psoriatic human skin spatial transcriptomics. **A**. Identified cell type clusters, **B-D**. Expressions of canonical cell type lineage marker genes for smooth muscle, eccrine gland, and fibroblast, **E**. Top ten spatially informed gradients generated from STew, **F-H**. Correlations between expressions of associated genes and specific sCVs. Pearson correlation and p-value with 95% confidence intervals are given for each test.

Next, we examined the identified low-dimensional components and found that these sCVs reflect different biologically meaningful spatial variations (**Fig. 5E**). For instance, sCV1 captures the gradual changing of expressions for keratinocyte-specific genes, including *SBSN, KRT2, FABP5* and *KRTDAP* (**Fig. 5F**). Similarly, sCV4 loadings are correlated with immune cell relevant gene expressions, in particular, myeloid cells (e.g., *LYZ*) and T cells (e.g., *CD4*) (**Fig. 5G**). Intriguingly, we also revealed that pro-inflammatory cytokines *CCL19* and *IL32*, detected by the statistical association from STew, are significantly associated with this axis, which further suggests an immune-derived spatial niche contributing to inflammatory disease pathogenesis (Pearson r = 0.3, *p* < 2.2e-16) (**Fig. 5G**). In parallel, sCV8 explains the smooth muscle cell axis (e.g., *ACTG2* and *DES*) (**Fig. 5H**), and sCV9 reflects potential fibroblast differentiation trajectory (e.g., *C3* and *SFRP2*) (**Fig. 5I**). In particular, *SFRP2* is an important marker for the pro-inflammatory fibroblast subpopulation discovered in psoriatic skin, which reflects the transition from a fibrotic to an inflammatory stage (Ma *et al*. 2023). In all, our derived low-dimensional embeddings review spatial niches that are associated with disease pathology.

## Discussion

Integrating spatial information into gene expression data is essential to understand the cellular activities in the tissue organization. In this study, we propose Stew, a method for identifying the shared information between spatial cell neighborhood relationships and the covariance observed in cell-cell similarities from the gene expression space. Our hypothesis is grounded in the notion that adjacent cells tend to exhibit a greater resemblance in transcriptional identities (Nitzan *et al*. 2019; Ren *et al*. 2020; Wang *et al*. 2023). Through rigorous evaluation and benchmarking with other state-of-the-art algorithms on four single-cell ST datasets, STew demonstrated superior performance, particularly in terms of clustering accuracy and delineation of smooth and continuous spatial domains. Furthermore, the representations derived from STew are effective in extracting biologically meaningful axes that reflect certain spatial niches, thereby facilitating the decryption of spatially informed cell neighborhood clusters. This progress marks a pivotal step towards a deeper understanding of the complex dynamics underlying cellular functions and tissue organization.

Although methods using deep learning-based models have been developed to decipher spatial domains by combining histological images with gene expression data (Pham *et al*. 2020; Zeng *et al*. 2023), several potential drawbacks exist. One notable shortcoming is that this may lead to poor prediction when the image features drive the construction of incorrect spot relationships, a problem exacerbated if the spot relations are not properly updated during the model training phase. Further, most of these models pass the crucial step of pre-training a specific big model on histological images for feature extraction. In addition, single-cell deep learning models frequently encounter issues with limited interpretability, and the embeddings they produce tend to be neither easily interpretable nor reproducible. In contrast, STew optimizes the covariance between gene expression patterns and spatial variations through a graph-embedded approach, which minimizes noise from individual data modalities and simultaneously learns representative features by enforcing sparsity constraints. This integrative strategy offers both scalability and interpretability, attributed to its linear characteristics, and the learned joint embeddings are reproducible given the optimization problem setting.

We note that the concept of using canonical correlation analysis has been used in genetics and microarray data integration (Parkhomenko, Tritchler and Beyene 2007, 2009), cross-dataset single-cell transcriptomics integration (Butler *et al*. 2018), and CITE-seq multi-modal data integration (Nathan *et al*. 2021; Zhang *et al*. 2022). STew effectively extends these ideas into a formal R package enriched with innovative functions specifically tailored to address spatial transcriptomics data challenges, including 1) embedding the data points in the graph-based representations, 2) automatically estimating the sparsity penalties to infer the influence of spatial correlation, 3) scalable and reproducible to handle large datasets, and 4) integrating a robust statistical goodness of fit method into STew to elucidate the gene signatures that hold significant associations with joint embeddings or spatially informed clusters. Altogether, STew is a comprehensive analytical R tool for single-cell ST datasets.

Further, STew can be extended and improved in multiple aspects. Given the potential in harnessing non-linear structures inherent in ST data (He *et al*. 2020; Biancalani *et al*. 2021; Zhao, Wang and Hu 2023), one prospective avenue we plan to explore is to extend this work by incorporating non-linear variations of canonical correlation analysis (Witten, Tibshirani and Hastie 2009), such as incorporating kernels or autoencoder structures, thereby fostering the learning of extracted features through non-linear processes. Considering the instances where cells in proximate spatial locales may not consistently present analogous transcriptional patterns–due to the dynamic interplay between tissue structures and cell type heterogeneity–we could add the “private variables” (Wang, Lee and Livescu 2016). These elements will help model unique variations from the spatial region and gene expression space, supplementing the already captured common variation. As the volume of spatial transcriptomics datasets continues to grow, we anticipate that STew will facilitate the discovery of new cellular organizations to help researchers identify new disease mechanisms.

## Code availability

The code and tutorials for STew package are available at https://github.com/fanzhanglab/STew.

## AUTHOR CONTRIBUTIONS STATEMENT

F.Z. conceived the study. N.G. implemented the algorithm and pipeline. J.V. implemented the statistical modeling. N.G., J.V, and D.F run all the benchmarking experiments, data analyses, and result visualizations. R.K. finalized the software package. N.G. and F.Z. wrote the initial draft. All authors participated in editing the final manuscript.

## Acknowledgments

We thank the Zhang Laboratory members for their valuable feedback and comments.

