## Supplementary information for "Uncover spatially informed shared variations for single-cell spatial transcriptomics with STew"

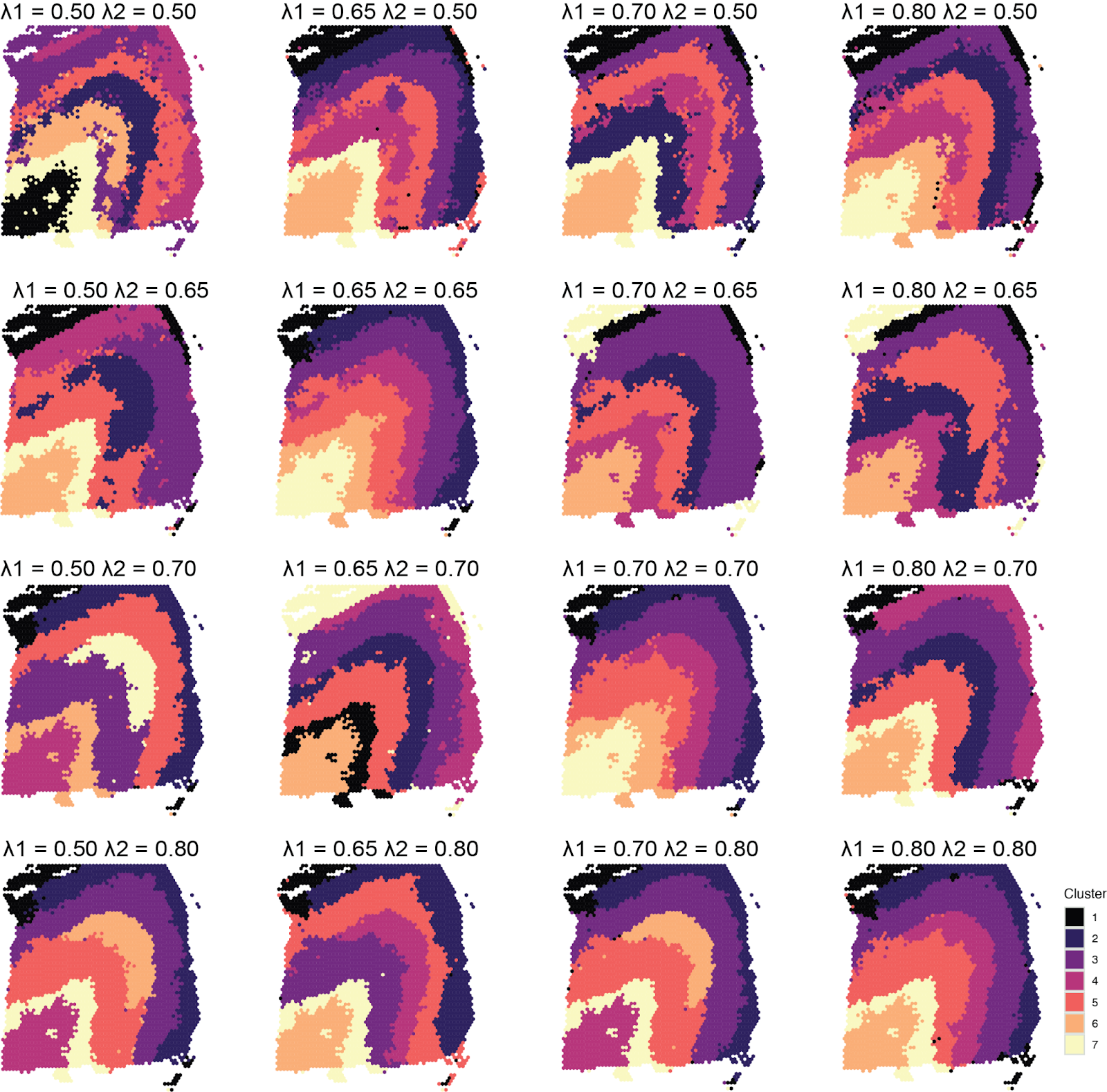

**Supplementary Figure 1.** Sensitivity analyses of running STew on the DLPFC data.  Identified cell type clusters from STew using different sparsity constraints. $\lambda_{1}$ denotes the penalty of gene expression-driven adjacency matrix. $\lambda_{2}$ denotes the penalty of spatial-driven cell neighborhood graph.

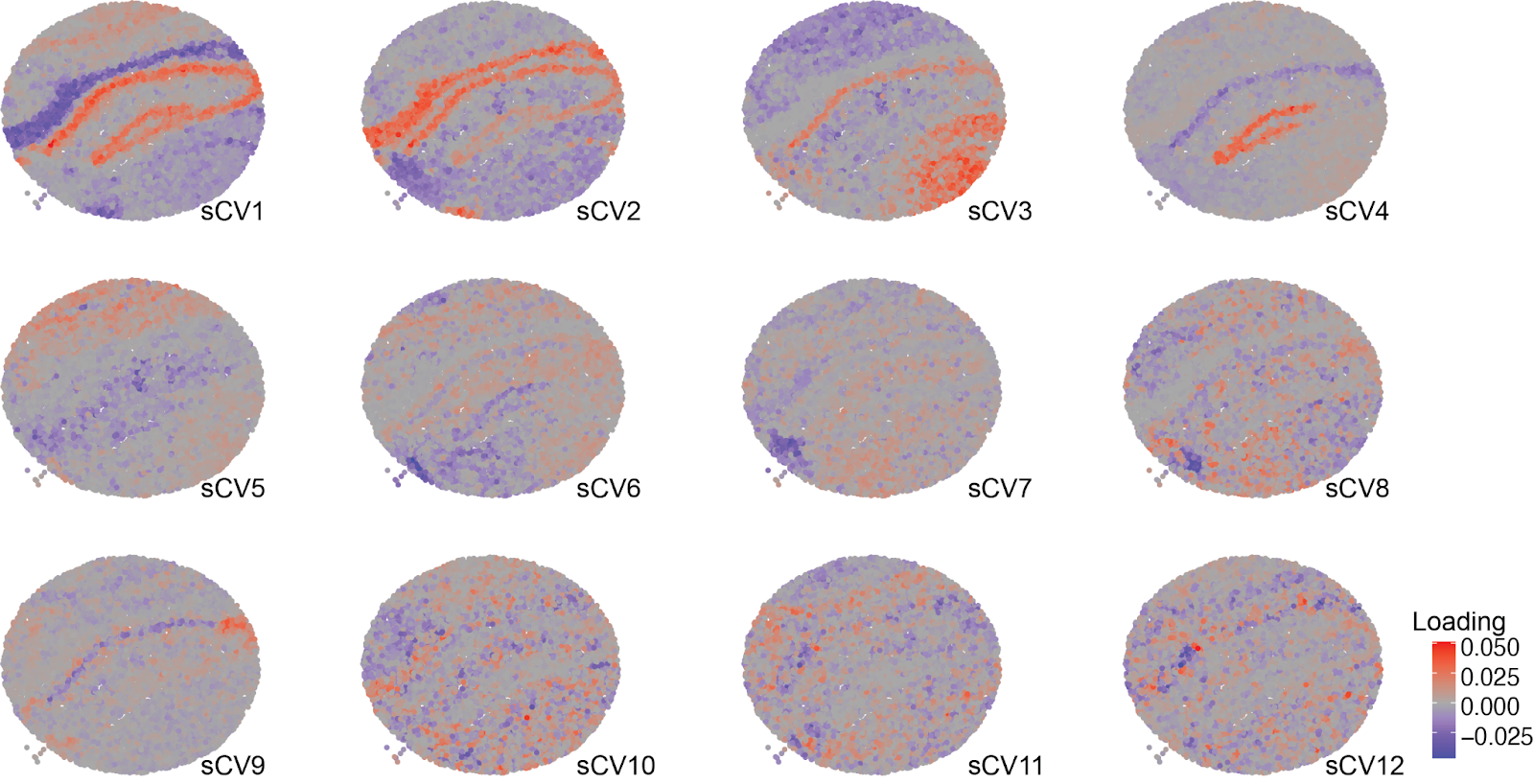

**Supplementary Figure 2.** Spatially informed gradients from STew on mouse hippocampus data from Slide-seqV2.

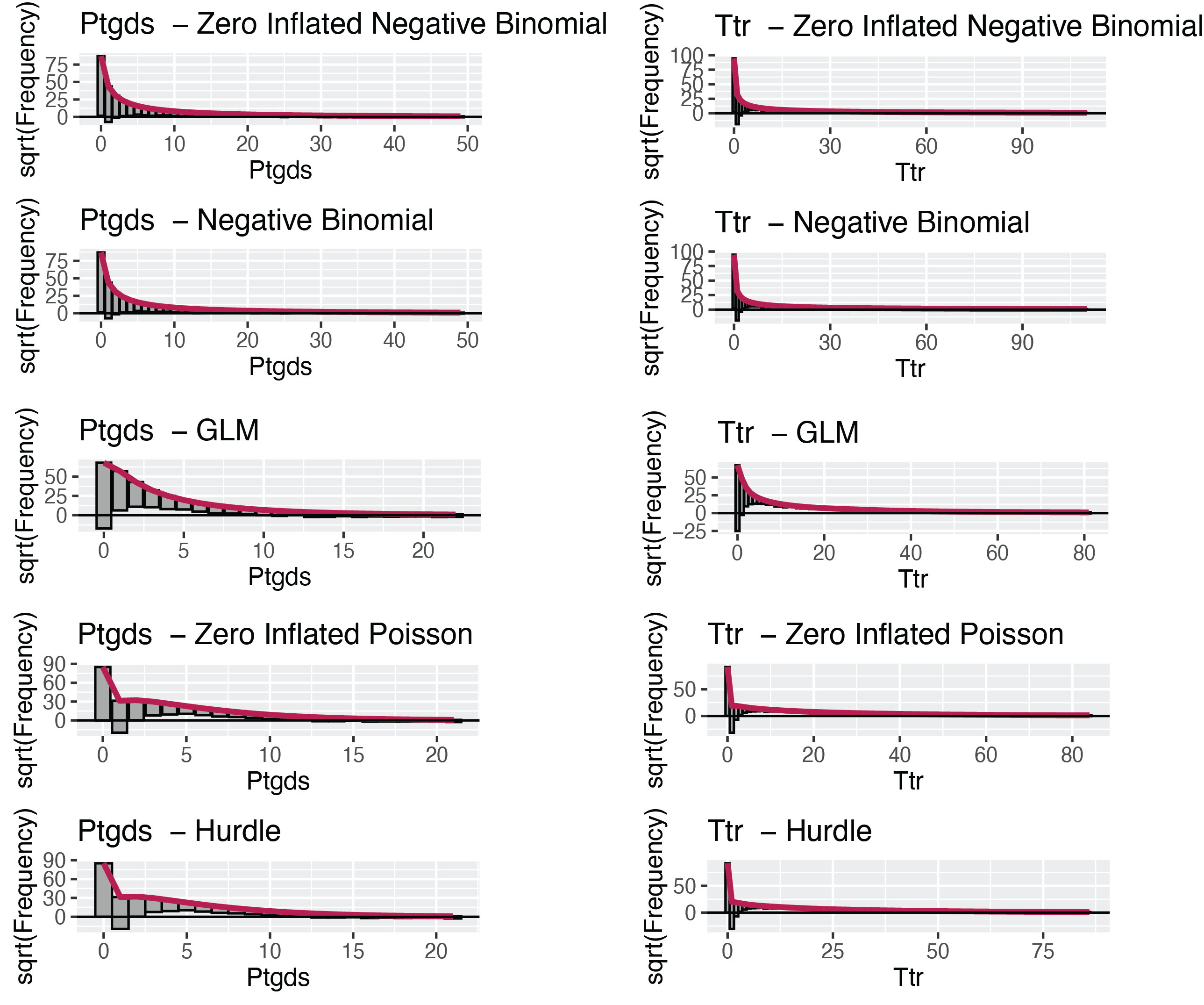

**Supplementary Figure 3.** Rootograms that measure squared residual frequency of model fits for genes *Ptgds* and *Ttr* from the Slide-seqV2 mouse hippocampus dataset.

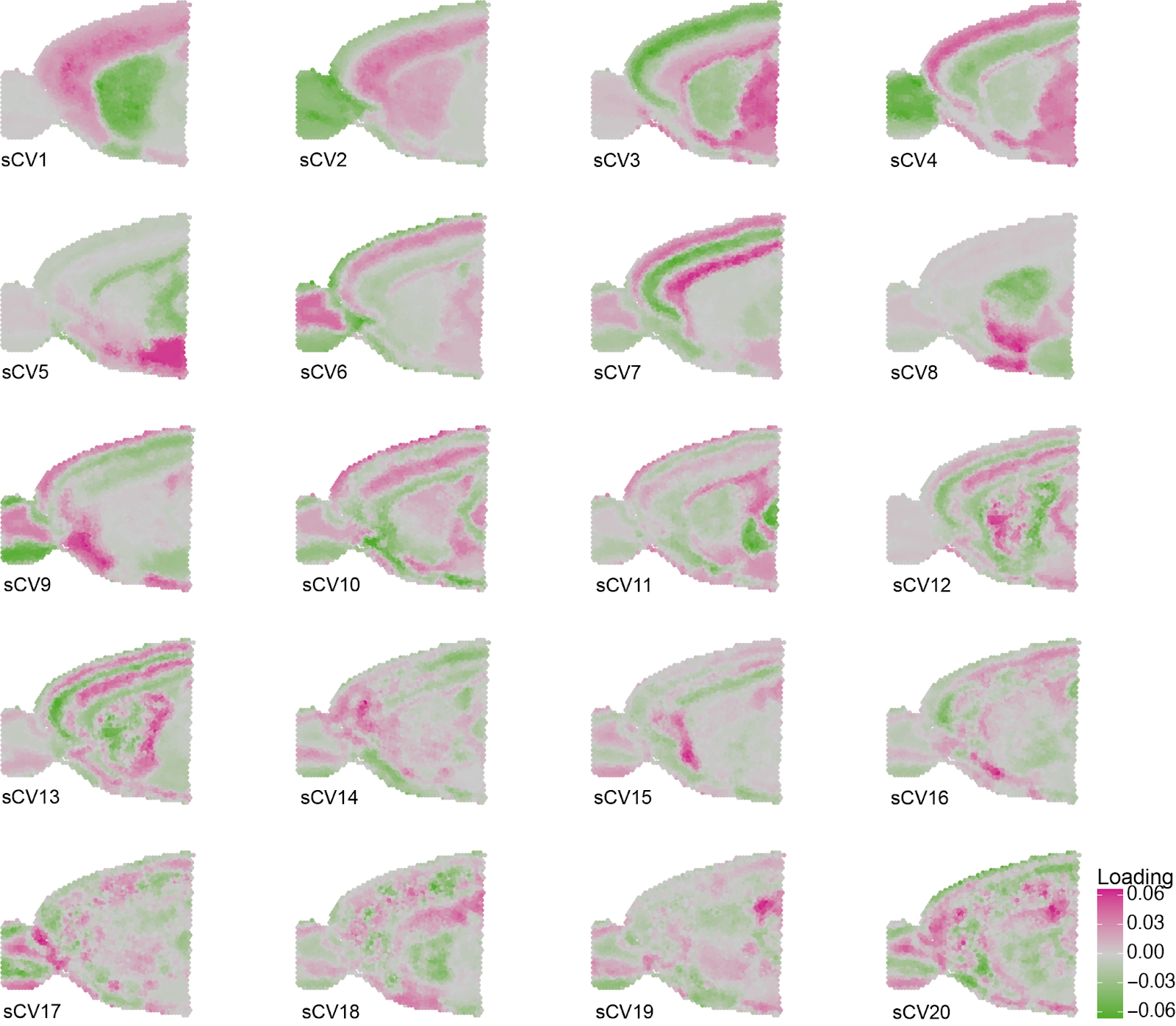

**Supplementary Figure 4.** Spatially informed gradients from STew on mouse brain anterior data from 10x Visium.
